# Improving human coronavirus OC43 (HCoV-OC43) research results comparability in studies using HCoV-OC43 as a surrogate for SARS-CoV-2

**DOI:** 10.1101/2021.07.14.452401

**Authors:** Erin E. Schirtzinger, Yunjeong Kim, A. Sally Davis

## Abstract

The severe acute respiratory syndrome coronavirus 2 (SARS-CoV-2) pandemic has renewed interest in human coronaviruses that cause the common cold, particularly as research with them at biosafety level (BSL)-2 avoids the added costs and biosafety concerns that accompany work with SARS-COV-2, BSL-3 research. One of these, human coronavirus OC43 (HCoV-OC43), is a well-matched surrogate for SARS-CoV-2 because it is also a *Betacoronavirus*, targets the human respiratory system, is transmitted via respiratory aerosols and droplets and is relatively resistant to disinfectants. Unfortunately, growth of HCoV-OC43 in the recommended human colon cancer (HRT-18) cells does not produce obvious cytopathic effect (CPE) and its titration in these cells requires expensive antibody-based detection. Consequently, multiple quantification approaches for HCoV-OC43 using alternative cell lines exist, which complicates comparison of research results. Hence, we investigated the basic growth parameters of HCoV-OC43 infection in three of these cell lines (HRT-18, human lung fibroblasts (MRC-5) and African green monkey kidney (Vero E6) cells) including the differential development of cytopathic effect (CPE) and explored reducing the cost, time and complexity of antibody-based detection assay. Multi-step growth curves were conducted in each cell type in triplicate at a multiplicity of infection of 0.1 with daily sampling for seven days. Samples were quantified by tissue culture infectious dose_50_(TCID_50_)/ml or plaque assay (cell line dependent) and additionally analyzed on the Sartorius Virus Counter 3100 (VC), which uses flow virometry to count the total number of intact virus particles in a sample. We improved the reproducibility of a previously described antibody-based detection based TCID_50_ assay by identifying commercial sources for antibodies, decreasing antibody concentrations and simplifying the detection process. The growth curves demonstrated that HCoV-O43 grown in MRC-5 cells reached a peak titer of ∼10^7^ plaque forming units/ml at two days post infection (dpi). In contrast, HCoV-OC43 grown on HRT-18 cells required six days to reach a peak titer of ∼10^6.5^ TCID_50_/ml. HCoV-OC43 produced CPE in Vero E6 cells but these growth curve samples failed to produce CPE in a plaque assay after four days. Analysis of the VC data in combination with plaque and TCID_50_ assays together revealed that the defective:infectious virion ratio of MRC-5 propagated HCoV-OC43 was less than 3:1 for 1-6 dpi while HCoV-OC43 propagated in HRT-18 cells varied from 41:1 at 1 dpi, to 329:4 at 4 dpi to 94:1 at 7 dpi. These results should enable better comparison of extant HCoV-OC43 study results and prompt further standardization efforts.

---

The severe acute respiratory syndrome coronavirus 2 (SARS-CoV-2) pandemic has renewed interest in human coronaviruses that cause the common cold, particularly as biosafety level (BSL)-2 surrogate viruses for SARS-CoV-2 research, which must be conducted at BSL-3 (Greenberg, 2016). Early in the current pandemic, the American Society for Testing and Materials (ASTM) issued guidance on SARS-CoV-2 surrogate virus selection for projects focusing on study of the persistence of the virus in different environments and the development and testing of decontamination procedures (ASTM 2020). The ASTM identified human coronavirus OC43 (HCoV-OC43) (order: Nidovirales, family: *Coronaviridae*, genus: *Betacoronavirus*, subgenus: *Embecovirus*, species: *Betacoronavirus 1*) as a preferred surrogate on account of its shared characteristics with SARS-CoV-2. As members of the same genus, *Betacoronavirus*, HCoV-OC43 and SARS-CoV-2 are closely related genetically (Lu et al., 2020). Both replicate in human respiratory epithelium and are transmitted by aerosols and droplets (Kutter et al., 2018). Although a rare occurrence, HCoV-OC43 has caused fatal encephalitis in immunocompromised humans. It can infect human neural cell lines and its neurotropic and neuroinvasive properties have been characterized in mice (Jacomy and Talbot, 2003; Jacomy and Talbot, 2006). Finally, HCoV-OC43 is more resistant than other human coronaviruses to quaternary ammonium disinfectants, such as benzalkonium chloride (Wood and Payne, 1998). Since the nature of our current research is focused accordingly, we selected HCoV-OC43 as our surrogate for methods development for our SARS-CoV-2 environmental virology research.

While using HCoV-OC43 as a surrogate for SARS-CoV-2 may have advantages over other coronaviruses or non-coronavirus surrogates, working with HCoV-OC43 also has its difficulties. Prior to the SARS-CoV and Middle East respiratory syndrome outbreaks in 2003 and 2012 respectively, research with human coronaviruses was fairly limited, likely because infections cause relatively mild disease, i.e. “the common cold,” and are typically restricted to the upper respiratory tract. Also, a review of extant literature finds researchers using a variety of methods to quantify HCoV-OC43 employing different cell lines, HRT-18 (human colon cancer cells) (Lambert et al., 2008), RD (human rhabdosarcoma cells) (Schmidt, Cooney, and Kenny, 1979), MRC-5 (human lung fibroblasts) (Collins, 1993; Collins, 1995; Kim et al., 2019), BSC-1(African green monkey kidney cells) (Bruckova et al., 1970), Vero E6 expressing TMPRSS2 (Hirose et al., 2021) and Mv1Lu (mink lung cells) (Bracci et al., 2020), using different titration assay types (either TCID_50_ or plaque assay), the latter with different types (agarose, Avicel, methylcellulose) and percentages of overlays, employing different incubation temperatures (33°C or 37°C), times (2-7 days) and detection methods (immunoperoxidase with chromagen, immunofluorescence, neutral red, crystal violet). This lack of standardization hampers evaluation and comparison of study results, as the different cell lines may have different growth kinetics and susceptibilities to virus grown in heterologous cells.

Therefore, in this study we sought to optimize and standardize existing HCoV-OC43 titration assays to facilitate comparison of study results from different researchers. Consequently, we optimized and standardized the indirect immunocytochemistry assay described by Lambert et al (2008) for quantification of HCoV-OC43 on HRT-18 cells. We examined the replication kinetics of HCoV-OC43 infection in the commonly used cell lines: HRT-18 (ATCC CCL-244), MRC-5 (ATCC CCL-171) and Vero E6 (ATCC CRL-1586) cells. We compared the susceptibility of HRT-18 cells to virus propagated on MRC-5 cells as well as vice versa. Finally, we measured the ratio of infectious to non-infectious or defective virions in virus stocks propagated in all three cell lines.

In order to increase the reproducibility of the antibody detection method of Lambert et al (2008), we identified commercial sources for the primary and secondary antibodies, mouse anti-HCoV-OC43 nucleoprotein (clone 542-7D) (Millipore Sigma) and donkey anti-mouse IgG Alexa Fluor 488 (Jackson Immunoresearch), respectively. In order to reduce the overall cost of the detection method, we first optimized the dilution of the primary antibody (see supplementary information for detailed methods). Briefly, 70-80% confluent HRT-18 cells in a 96-well plate format were infected with 10-fold serial dilutions of HCoV-OC43 low passage stock virus and incubated at 33°C, 5% CO_2_ for four days. The cell supernatant was removed, the cells washed with 1X PBS and 4% paraformaldehyde added to fix the cells for 30 minutes. The cells were incubated for 2 hours at 37°C with multiple primary antibody dilutions in 1X PBS including 1:500 (manufacturer’s recommended dilution), 1:1000, 1:2000, 1:4000 and 1:10000. Post three 1X PBS washes, the cells were incubated for 2 hours in the dark at 37°C with secondary antibody diluted 1:200 in 1X PBS (the high end of manufacturer’s recommended range). Post wash, 1X PBS was added to the wells and the plates were read on an Olympus CKX53 epifluorescence microscope using a 10X/0.25 NA objective. The highest dilution of primary antibody that showed no signal loss in comparison with lower dilutions was chosen as the optimal dilution. After selecting the primary antibody dilution, the secondary antibody was titrated similarly using dilutions of 1:200, 1:400 and 1:800. Finally, we optimized the antibody volumes by comparing 100 µl to 50 µl per well, secondary antibody incubation time by comparing the signal from a two-hour incubation to a 30-minute incubation (manufacturer’s recommendation) and decreased the paraformaldehyde fixation time from 30 to 10 minutes.

We investigated the replication kinetics of HCoV-OC43 in HRT-18 cells in comparison to Vero E6 and MRC-5 cells by conducting multi-step growth curves in each cell type. In triplicate, 90% confluent T25 flasks were infected with HCoV-OC43 at a multiplicity of infection of 0.1 in cell type matched media, Roswell Park Memorial Institute (RPMI) 1640, ATCC formulation (Gibco, Fisher Scientific) supplemented with 2% heat-inactivated fetal bovine serum (HI-FBS) (Gibco, Fisher Scientific), 2mM L-glutamine (Corning, Fisher Scientific) and 1X Penicillin/Streptomycin (Corning, Fisher Scientific) for the HRT-18 cells and Eagle’s Minimum Essential Medium (Corning, Fisher Scientific) supplemented with 2% HI-FBS and 1X Penicillin/Streptomycin (Corning, Fisher Scientific) for the Vero E6 and MRC-5 cells. The flasks were incubated at 33°C, 5% CO_2_ for 1.5 hours with rocking every 15 minutes. The inoculum was removed, the flasks washed twice with 1X PBS and 5 ml of media added to each flask. One ml of supernatant was collected and one ml of fresh media added every 24 hours for seven days. The samples were aliquoted and stored at -80°C until assayed by TCID_50_ or plaque assay as well as flow virometry. The development of cytopathic effect (CPE) was monitored using phase contrast (4X/0.13 NA and 10X objectives) on the aforementioned microscope every 24 hours, immediately prior to sample collection. Once CPE developed in a cell type, a representative flask was photographed daily through day 7 at 40x and 100x magnification using an Olympus LC30 camera and cellSens Entry software version 1.16.

For the HRT-18 experiment, the samples were quantified by TCID_50_ on HRT-18 cells as described above. For the MRC-5 and Vero E6 experiments, samples were quantified by plaque assay. Briefly, monolayers of MRC-5 or Vero E6 cells in 12- or 24-well plates were inoculated with 10-fold serial dilutions of virus sample in triplicate (12-well) or duplicate (24 well). The inoculum was adsorbed for 1.5 hours at 33°C and 5% CO_2_ with rocking every 15 minutes. The inoculum was removed and 1% methylcellulose in EMEM with 2% HI-FBS and 1X P/S overlay applied. The plates were incubated at 33°C and 5% CO_2_ for four days after which the inoculum was removed and the cells incubated in crystal violet fixative (25% v/v of 37% formaldehyde, 11% v/v of 95% ethanol, 5% v/v of glacial acetic acid and 4% w/v crystal violet in deionized water) for one hour at room temperature. The crystal violet fixative was removed, plates washed with tap water, plaques counted and titers calculated.

The Virus Counter 3100 (Sartorius Stedim) was used to determine the total number of virus particles (infectious and non-infectious) by flow virometry in all the HCoV-OC43 samples collected during the propagation on the three cell lines. Prior to sample titration, a screening assay was conducted, using the Combo Dye kit (Sartorius) per manufacturer’s instructions, to determine the appropriate dilution for the virus samples. Briefly, Combo Dye Concentrate was rehydrated and diluted with Combo Dye Buffer. The cell culture media (used as a control for background signal in the media) and the HCoV-OC43 stock virus were diluted at 1:10, 1:100, 1:1000 and 1:10000 in Sample Dilution Buffer, 100 µl of each dilution and 50 µl of the diluted Combo Dye were added to each sample vial, mixed and incubated at room temperature in the dark for 30 minutes. The samples were then run on the Virus Counter and the optimal dilution range was calculated using the Virus Counter software. Once the appropriate dilution was determined, all samples from each of the growth curves were diluted, stained with Combo Dye as described above and run on the Virus Counter in duplicate. The average titers of all sample duplicates were calculated using the Virus Counter software. For a titer to be considered reliable the data acquired for both run needs to exceed two internal quality thresholds.

The Lambert et al (2008) immunoperoxidase detection based TCID_50_ assay has not been widely adopted. The detection protocol is complex, time-consuming (> 5 hours) and expensive. Consequently, we identified commercial sources of antibodies to increase the reproducibility of the assay and optimized the antibody dilutions to reduce the cost. We found 1:2000 and 1:400 dilutions to be optimal for primary and secondary antibodies, respectively. We also found that reduction of the antibody volume from 100 µl to 50 µl per well in a 96 well plate format resulted in no signal loss and halved antibody volume required per assay. Overall, these changes reduced the antibody cost per assay by 67%. Additionally, we reduced the total detection protocol time to approximately three hours by decreasing the secondary antibody incubation time to 30 minutes with no signal loss, decreasing the cell fixation time to 10 minutes, and eliminating the reporter enzyme and chromogen steps through use of a fluorophore conjugated secondary antibody.

This investigation of HCOV-OC43 replication kinetics in HRT-18, MRC-5 and Vero E6 cells provided several insights regarding the use of MRC-5 and Vero E6 cells in HCoV-OC43 quantification. As expected, HRT-18 cells infected with HCoV-OC43 showed no obvious CPE at any time point. In contrast, HCoV-OC43 produced CPE (cell lysis, cell rounding and vacuolization) in MRC-5 cells starting on day 2 which continued through day 7. Also, contrary to some previous reports (Bracci et al 2020; Hirose et al 2021), CPE was observed in HCoV-OC43 infected Vero E6 cells. Cell rounding and lysis began on day 3 and continued until day 7 (Fig. 1-2).

**Figure 1.**
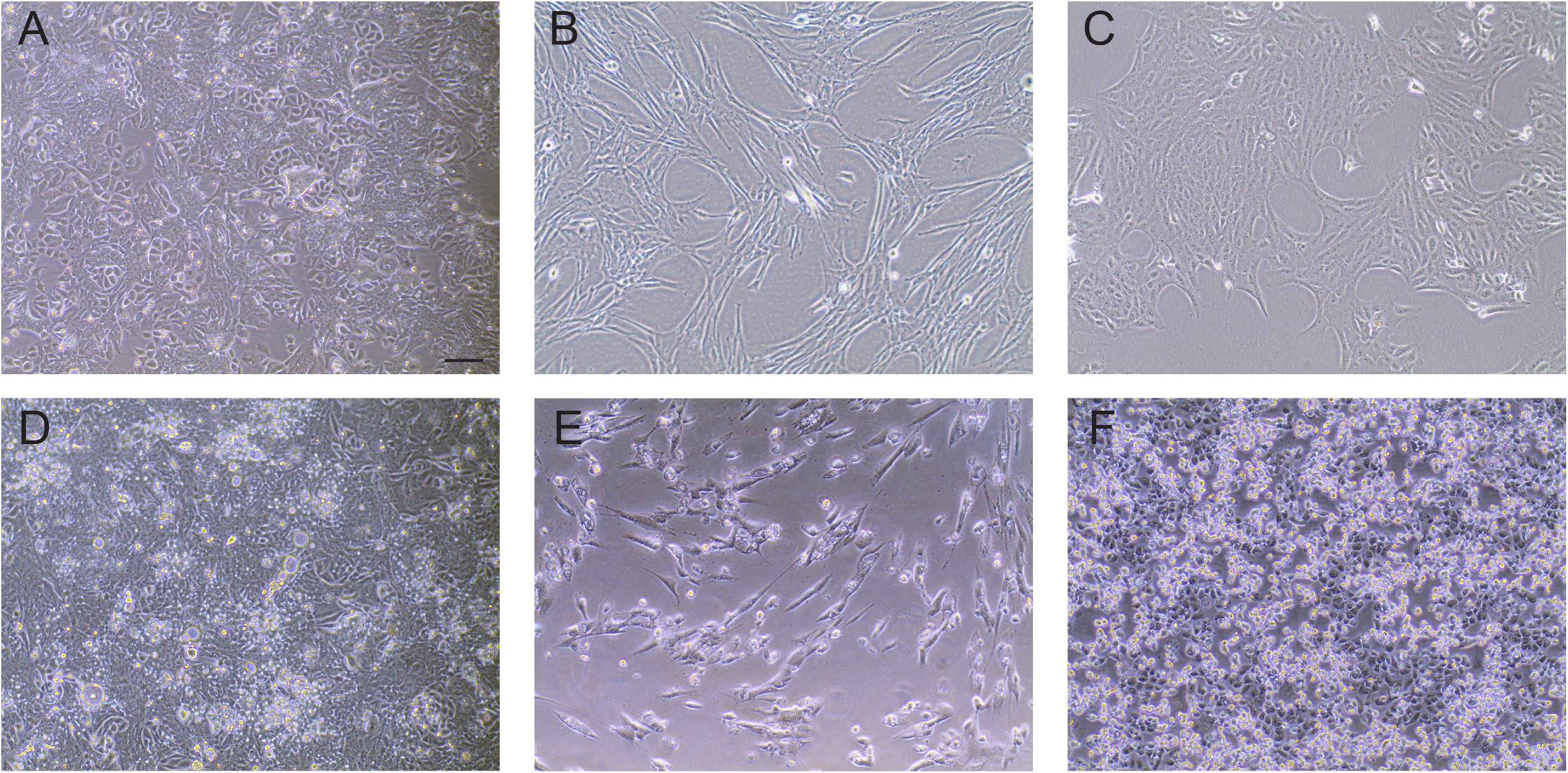
Comparison of uninfected and HCoV-OC43 infected cells. Uninfected cells: A) HRT-18, human ileocecal colorectal adenocarcinoma (ATCC CCL-224), B) MRC-5, human lung fibroblast (ATCC CCL-171), C) Vero E6, African green monkey kidney cells (clone E6 from Vero 76) (ATCC CCL-1586). Cells infected with HCoV-OC43 at four days post infection (dpi): D) HRT-18, E) MRC-5, F) Vero E6 cells. All images were taken on an Olympus CKX35 microscope equipped as described in the methods. Bar is 100 microns.

**Figure 2.**
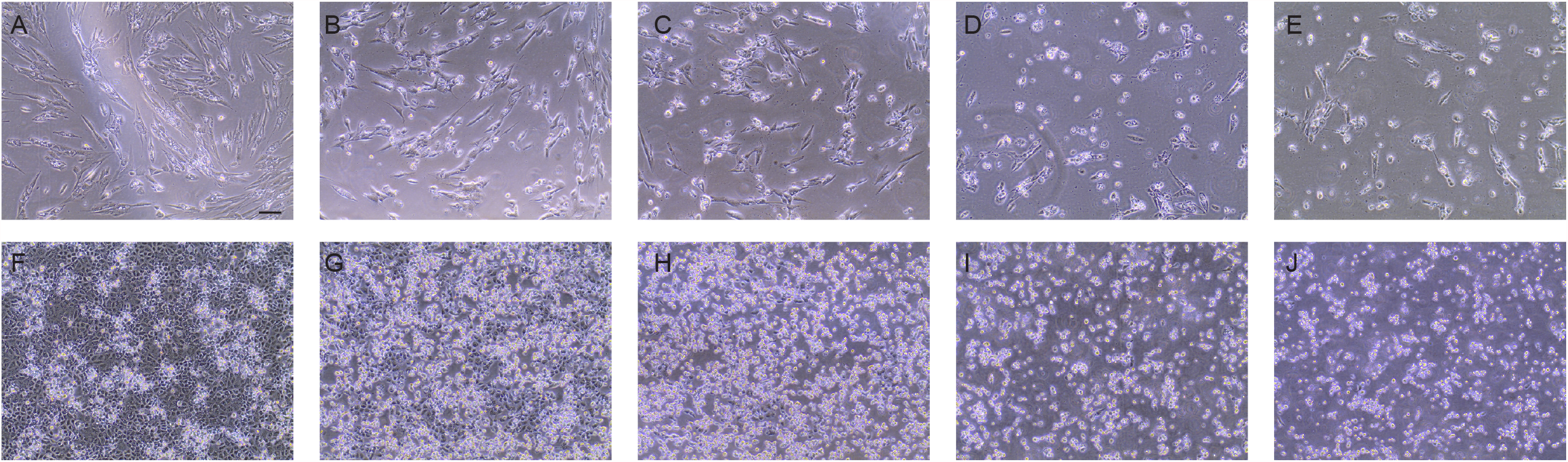
Cytopathic effect in HCoV-OC43 infected cells over time. MRC-5 cells at (A-E) 3-7 dpi, one image per day; (B) is same image as in Fig. 1e. Vero E6 cells at (F-J) 3 -7 dpi, one image per day; (G) is same image as Fig. 1f. Bar is 100 microns.

Plaques produced by HCoV-OC43 infection of the MRC-5 cell monolayers were enumerable at 40x magnification on a phase contrast microscope. However, the morphology of the cells made oddly shaped plaques (Fig. 3a-c) that quickly grew into each other at lower dilutions during the four-day incubation. Bracci et al (2020) tested HCoV-OC43 in a plaque assay using MRC-5 cells with a five-day incubation at 37°C. They concluded that plaques were countable, but that MRC-5 cells were too slow-growing and variable to further optimize the assay (Bracci et al 2020). The incubation time for plaques to develop in MRC-5 cells may need to be shortened to less than four days to prevent the plaques from growing together or a higher percentage overlay used such that its viscosity better restricts the movement of the virus. Previous studies that used MRC-5 cells for HCoV-OC43 plaque assays have used agarose overlays, which may better limit the spreading of plaques (Collins 1995; Funk et al 2012; Warnes et al 2015). Alternatively, using MRC-5 cells in a TCID_50_ format for HCoV-OC43 quantification would alleviate the need for distinguishable plaques and might be a useful assay when greater throughput is needed.

**Figure 3.**
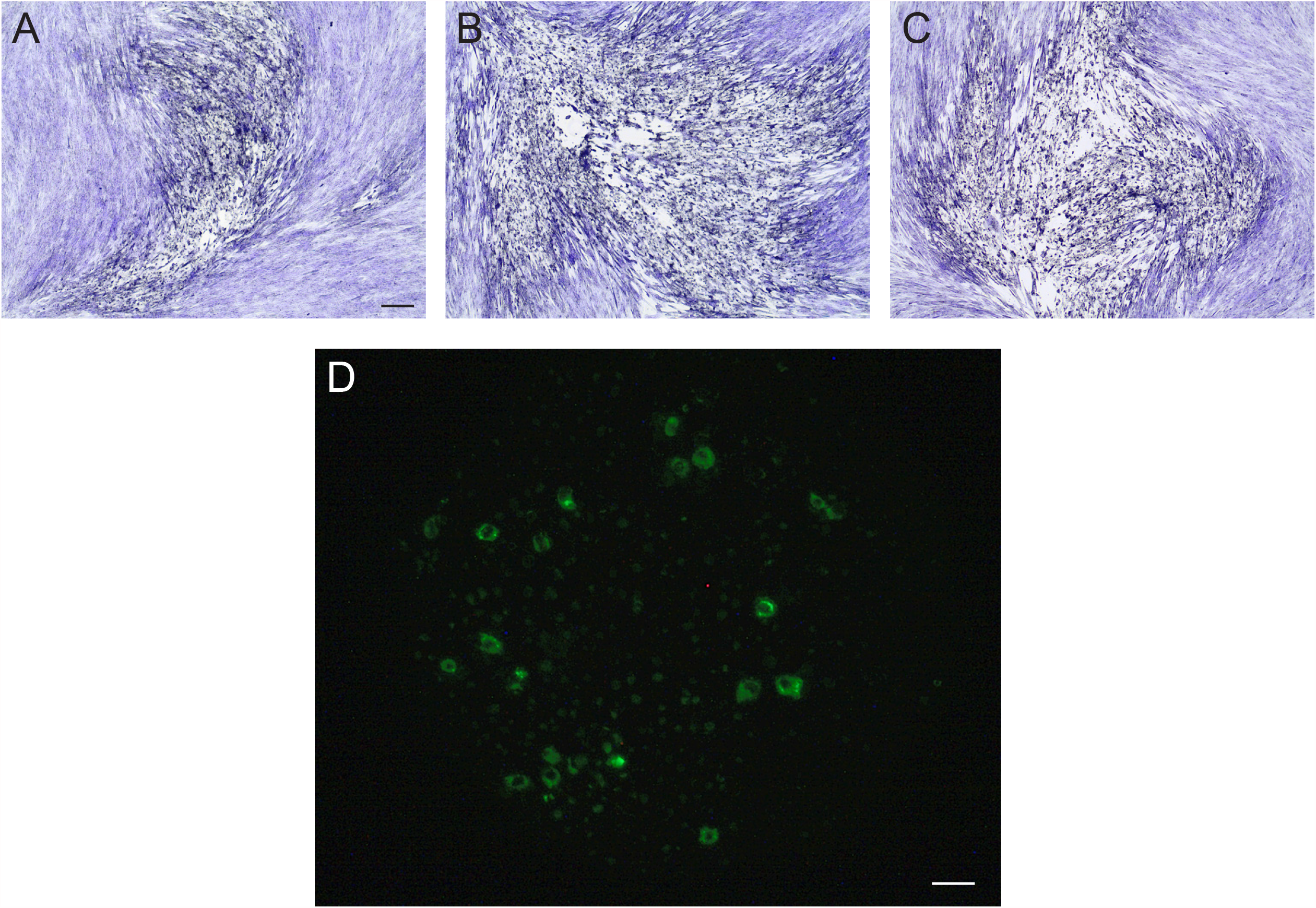
Visualization of HCoV-OC43 infection of different cell lines. Detection of HCoV-OC43 infection using the optimized immunofluorescence assay: ((A-C) Crystal violet staining of plaques in MRC-5 cells, showing pleomorphic plaque morphology. D) Vero E6 cells. The black bar is 100 microns and applies to (A-C). The white bar is 20 microns and applies to (D).

In contrast, after a four-day incubation, there were no plaques visible in the Vero E6 propagated HCoV-OC43 plaque assays using Vero E6 cells. This result was unexpected due to the CPE observed during propagation. To determine if Vero E6 cells could be used for quantification of HCoV-OC43 propagated on other cell lines, HRT-18 cell-propagated HCoV-OC43 virus was tested in a TCID_50_ assay using Vero E6 cells. At 7 days of incubation, the cells were fixed and virus detected by immunofluorescence as described above. Immunofluorescence confirmed the HCoV-OC43 infection up to the 10^−5^ dilution in the Vero E6 cells (Fig. 3d), however CPE was only visible in the lowest (10^−1^) dilution on the plate. The Vero E6 growth curve samples were then titrated by TCID_50_ on HRT-18 cells as described above to determine the virus’s basic growth parameters in the Vero E6 cell line. When Hirose et al (2021) similarly tested a Vero E6 plaque assay, they found that infection of Vero E6 cells with 2 × 10^5^ TCID_50_/ml of HCoV-OC43 produced barely visible CPE from 3 to 5 days post-infection (dpi) while infecting Vero E6 cells with less than 2 × 10^3^ TCID_50_/ml of HCoV-OC43 resulted in no CPE during a 5-day infection period. The lack of CPE in the current experiment as well as in Hirose et al (2021) may have been caused by terminating the incubation period prior to the virus reaching sufficient levels to cause CPE. The length of time needed for low titer HCoV-OC43 to develop CPE was not determined in our study, but based on our data it is greater than 7 days. This extended incubation time needed for HCoV-OC43 to produce CPE in Vero E6 cells precludes their usefulness in a standard method of quantification and emphasizes the need to know the susceptibility of the cell line being used for quantification.

The growth curves of HCoV-OC43 in HRT-18 and Vero E6 cells shared several similarities (Fig. 4a and b). At 1 dpi the log_10_ TCID_50_/ml values were approximately 5.0 and similar peak titers of approximately 6.5 log_10_ TCID_50_/ml were reached at 5 and 6 dpi in Vero E6 and HRT-18 cells, respectively. Due to the nature of TCID_50_ assays being indirect endpoint assessments of the number of infectious virions, these numbers over estimate of infectious titer. In contrast, the MRC-5 cell-based titration by plaque assay is more accurate because each well-separated plaque directly represents a single infectious virion and because the overlay restricts its progeny virus infecting neighboring cells. Curiously, the HCoV-OC43 virus grown in MRC-5 cells is nearly 2 log_10_ higher at 1 dpi than the TCID_50_/ml titers of the HRT-18 and Vero E6 cells (Fig. 4c). In addition, the MRC-5 virus reaches peak titer of 6.9 log_10_ plaque forming units (PFU)/ml which is approximately 1 log_10_ higher than the peak titer of HCoV-OC43 grown on HRT-18 cells at 6 dpi and approximately 0.5 log_10_ higher than HCoV-OC43 grown on Vero E6 cells at 5 dpi. The HCoV-OC43 reaches a higher titer, sooner, when grown in MRC-5 cells than the more commonly used HRT-18 cells.

**Figure 4.**
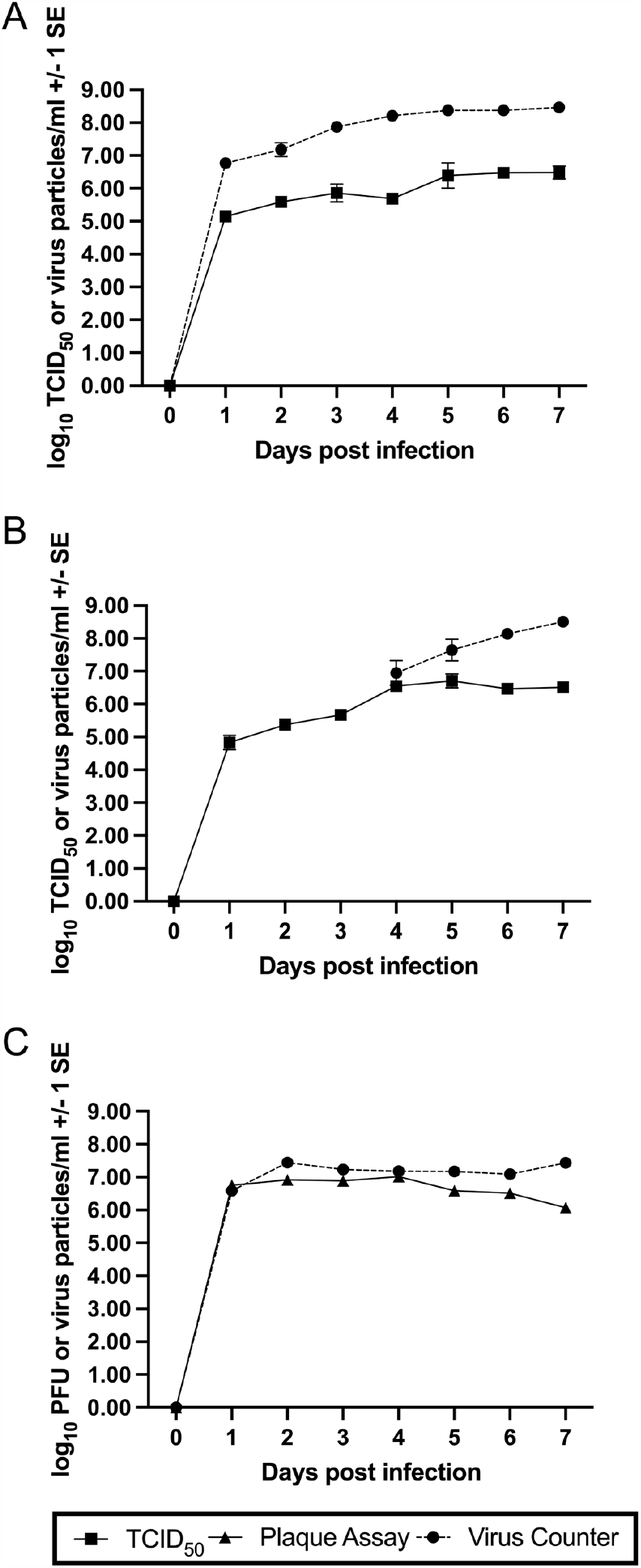
Comparison of infectious and total (infectious and defective) virions produced during HCoV-OC43 propagation of different cell lines over seven days. Closed squares and solid lines are infectious titers determined by TCID_50_ assay. Closed triangles and solid lines are infectious titers determined by plaque assay. Closed circles and dashed lines are total viral particles determined by Virus Counter analysis. All values are shown +/- 1 standard error. HCoV-OC43 infection of (A) HRT-18, (B) Vero E6 and (C) MRC-5 cells.

The cell lines used in this study were also differentially permissive to HCoV-OC43 virus grown in other cells. HCoV-OC43 grown in MRC-5 cells yielded a 7.57 log_10_TCID_50_/ml titer in MRC-5 cells but when assayed in HRT-18 cells the titer was greater than 10.5 log_10_TCID_50_/ml. HCoV-OC43 grown in Vero E6 cells developed no plaques despite showing CPE during the replication kinetics experiment. However, we were able to successfully quantify these samples using the TCID_50_ assay with HRT-18 cells. This suggests that HCoV-OC43 may use a different mechanism to enter Vero E6 cells and fewer cells may become infected per unit time. Further support for this idea came from a TCID_50_ assay of the stock HCoV-OC43 virus (grown in HRT-18 cells) using Vero E6 cells in which CPE was present only in the 10^−1^ dilution at 7 dpi but an HRT-18 based log_10_TCID_50_/ml titer was 6.0. Understanding that virus titer can change based on which cell lines are used to propagate as well as quantify the virus is important when comparing results from multiple studies, especially in a field with few gold standards for assays. Based on our investigations, we would recommend propagating virus and quantifying samples on the same cell type.

The Virus Counter 3100 (Sartorius Stedim) was used to determine the total virus particles (infectious and non-infectious) produced by HCoV-OC43 in each cell line. This instrument uses the principles of flow cytometry to count intact virus particles. Proprietary dyes separately label the nucleic acid and envelope proteins of a virion. As the sample passes through a capillary a laser excites the dyes and a count is made of nucleic acid, protein or both. Simultaneous detection of both dyes by the paired photomultiplier tubes indicates the presence of an intact enveloped virus particle. The titer of the sample is then calculated from the virus particle counts. Although, the Virus Counter 3100 greatly reduces the time needed to quantify virus samples, it is unable to determine if a virus particle is infectious. Consequently, use of the Virus Counter in combination with assays that determine infectious virus concentration, as done here, enabled determination of the number of defective virus particles (defective virions) produced at different points during propagation. Defective virions are produced when a mutation or deletion occurs during replication or a mistake is made in virus packaging (Alnaji and Brooke, 2020; Vignuzzi and Lopez, 2019). Often defective virions are able to enter cells, at times via alternative paths such as macropinocytosis, but they fail to complete the replication cycle without a co-infecting helper virus. When a helper virus is present the defective genome may have an advantage as the smaller defective genome can be replicated faster. In addition, competition for resources within the cell can negatively impact the production of infectious virus, driving down viral titers (Alnaji and Brooke, 2020; Makino, Taguchi, and Fujiwara, 1984). Defective virions are known to impact viral persistence, pathogenesis and evolution (Vignuzzi and Lopez, 2019). Therefore, it is important to monitor the production of defective virions in virus stocks to prevent unusual and ungeneralizable results.

Figure 4 shows the virus particle/ml titers from Virus Counter analysis (black circles, dotted lines) of HCoV-OC43 grown in each cell line over time. HCoV-OC43 grown on HRT-18 cells produced 6.77 log_10_ viral particles/ml at 1 dpi. The number of viral particles steadily increased to 8.46 log_10_ viral particles by 7 dpi. Subtracting the number of infectious virions from the number of total viral particles results in the number of defective virions. Dividing the number of defective virions by the number of infectious virions gives the ratio of defective to infectious virions (D:I) (Table 1S). At 1 dpi, HCoV-OC43 grown on HRT-18 cells has a the D:I of 41.1:1. The ratio peaks at 329.4:1 at 4 dpi and then decreases to 94.3:1 at 7 dpi. (Table 1S). In contrast, for the HCoV-OC43 grown in Vero E6 cells, however, the number of total virus particles did not surpass the Virus Counter’s minimum threshold (5.5 × 10^5^ particles/ml) until 4 dpi on which the log_10_ virus particles/ml was 6.95. The concentration of virus particles/ml continued to increase until 7 dpi when total virus particles were nearly 100 times the TCID_50_/ml. This pattern suggests that in the early part of the growth curve the less fit HCoV-OC43 genomes in Vero E6 cells are selected against because they fail to replicate. Defective virions may then increase after a threshold of infection wherein co-infections can occur. For our experiment, this threshold appears to occur between 3 dpi and 4 dpi when the log_10_ TCID_50_/ml is between 5.67 and 6.55 and the D:I at 4 dpi is 1.5.

Interestingly, for the HCoV-OC43 grown in MRC-5 cells the number of virus particles/ml was very similar (within <1.0 log_10_) to the number of PFU/ml throughout most of the sampling period (Fig. 4b). The D:I was 2.4:1 at 2 dpi, dropped to 0.5:1 at 4 dpi and peaked at 22.1:1 at 7 dpi. Based upon these preliminary results, HCoV-OC43 grown in MRC-5 cells may be a better choice for virus production due to the limited number of defective virions compared to HCoV-OC43 grown in HRT-18 cells. However, we did not continue to passage HCoV-OC43 in MRC-5 cells so it is possible that defective virions may increase with further passaging. The impact of the immune response of MRC-5 cells to HCoV-OC43 infection and the production of defective virions is also currently unknown.

In summary, our study showed that several protocols for quantification of HCoV-OC43 could be optimized and further standardized to encourage their wider use among researchers. We improved the indirect immunoperoxidase detection method of Lambert et al (2008) by identifying commercial sources of antibodies and reducing the time, cost and complexity of the protocol by adjusting incubation times in multiple steps, optimizing antibody dilution and volume and changing the detection system. Our comparison of growth parameters of HCoV-OC43 propagation in three cell lines (HRT-18, MRC-5 and Vero E6) found that HCoV-OC43 propagated in MRC-5 cells achieved a higher titer (∼7.0 log_10_ PFU/ml) that peaked at 2 dpi compared to that in HRT-18 cells (6.5 log_10_ TCID_50_/ml at 6 dpi). Our findings showed that Vero E6 cells are susceptible to HCoV-OC43 infection and infection at an MOI of 0.1 results in CPE. Finally, we calculated the D:I for HCoV-OC43 grown in each cell line. The virus propagated in MRC-5 cells had the lowest D:I, ranging from 0.5:1 to 2.9:1 over 6 days, then rising to 22.1:1 at 7 dpi. HCoV-OC43 propagated on HRT-18 cells had the highest D:I of 329.4:1 on 4 dpi, while the Vero E6 propagated HCoV-OC43 had a D:I of 1.5:1 at 4 dpi. The HRT-18 and Vero E6 propagated viruses, respectively, had ratios of 94:1 and 97:1 at 7 dpi. We hope that our findings will improve the consistency and reproducibility of HCoV-OC43 research, particularly when this virus is used as a surrogate for SARS-CoV-2.

